# Multi-Omics Characterization of Human Molecular Responses to Spaceflight Across Two Independent Missions

**DOI:** 10.64898/2026.06.01.729304

**Authors:** Abirami Santhanam, Zeineen Momin, Xiang Qin, Qiaoyan Wang, Aparna Krishnavajhala, Qi Jiang, Kimberly Walker, Divya Kalra, Marie-Claude Gingras, Hsu Chao, Kavya Kottapalli, Sravya V. Bhamidipati, Mishfak A.M. Mansoor, Sarah A. Ashiqueali, S. Michelle Griffin, Michal M. Masternak, Jimmy Wu, Donna Muzny, Emmanuel Urquieta, Richard A. Gibbs, Harsha Doddapaneni

## Abstract

Human spaceflight has historically been led by government agencies, but the emergence of commercial organizations is enabling broader participation and new research opportunities. In this study, we present a comprehensive molecular characterization of early human responses to spaceflight, leveraging multi-omics data across the first three weeks of two commercial missions. Biospecimens from six individuals, four from the Axiom 2 mission (10 days) and two from Axiom 3 (21 days), were analyzed using single-cell and bulk RNA sequencing, alongside proteomic profiling. Individual and integrative analyses of these datasets reveal systemic changes in cell types, transcripts, and proteins related to immune regulation, osteoclast differentiation, NF-κB signaling, and blood homeostasis pathways. Importantly, several of the detected pathways align with physiological patterns observed in longer-duration missions. This work establishes a foundational resource for understanding early adaptation to spaceflight at the cellular and molecular levels, providing insights to reduce future space-travel health risks.

**Highlights:**

- First integrated multi-omics analysis using single-cell, bulk RNA sequencing and proteomic profiling of early human spaceflight responses across two commercial missions (Ax-2 and Ax-3).
- Distinct PBMC clustering was observed across Ax-2 and Ax-3, and post-flight samples. It showed fewer monocytes, dendritic cells, and megakaryocytes with increased naïve CD4⁺ and cytotoxic T cells.
- Multi-omics integration identifies shared biological signatures, including osteoclast differentiation, metabolic stress, and coagulation changes.
- These findings lay the foundation for developing countermeasures to protect immune, skeletal, and vascular health during spaceflight.

## Introduction

Human space travel stands at a critical frontier for scientific discovery, human health, and survival. Advancing human space exploration is imperative to secure our future and push the boundaries of knowledge and innovation. Spaceflight, however, presents a complex array of physiological challenges due to exposure to microgravity, ionizing radiation, isolation, and other perils. These stressors disrupt homeostasis and trigger systemic responses, including immune dysregulation, musculoskeletal degradation, and cardiovascular remodeling. These disturbances lead to muscle atrophy, bone loss, and increased infection risk ^1,2^.

While sub-orbital missions of a few days, weeks, or up to a year on ISS have become routine and are deemed relatively safe for humans, the impact on the human body remains understudied, especially at the cellular and molecular level. Consequently, future long-duration missions demand a comprehensive understanding of molecular and physiological adaptations during short-term missions. This knowledge will be essential to safeguard astronaut health and ensure the success of these endeavors^3^ (https://www.nasa.gov/humans-in-space).

Astronaut studies are often constrained by small sample sizes or the use of single assays, limiting statistical power. To address these limitations, multi-omics approaches offer a framework to integrate diverse datasets for examining molecular and physiological changes. Therefore, there is a growing interest in applying omics technologies to study these effects. Such insights are critical for safeguarding astronaut health and ensuring mission success.

Two previous studies have applied multi-omics to study the physiology of space travel, the first being NASA’s Twins Study, which examined identical twin astronauts during a one-year mission^4^, and more recently, the Inspiration4 mission - the first all-civilian crew in low Earth orbit, for three days^5^. The GENESTAR (at Baylor College of Medicine-Human Genome Sequencing Center) initiative has advanced human spaceflight biology research by providing a unified framework for Clinical Laboratory Improvement Amendments (CLIA) grade standardized biospecimen collections for CLIA and research use, along with multi-omics analysis across missions and individuals^6^.

For the first time, this study provides a comprehensive molecular perspective on the human response during the first three weeks of the human response in spaceflight. A multi-omics dataset of single-cell RNA-seq, RNA-Seq (bulk), and proteomics from a multinational crew of six individuals, four from Axiom 2 (10 days) and two from Axiom 3 (21 days), is presented here. Together, these data provide a high-resolution resource for characterizing early spaceflight-associated molecular changes across commercial missions.

Studying such short-duration missions is crucial for identifying biological analogues relevant to long-term spaceflight. Physiological changes noted from prior studies in early weeks of spaceflight, such as internal jugular vein thrombosis (IJVT)^7^, spaceflight-associated neuro-ocular syndrome (SANS)^8^, immune dysregulation, and vascular or coagulation shifts ^9^, also highlight how rapidly the body responds to the space environment. By capturing these initial adaptations, molecular data from missions like Axiom 2 (Ax-2) and Axiom 3(Ax-3) serve as valuable testbeds, bridging Earth-based simulations and deep-space realities, and enabling the development of targeted, data-driven countermeasures.

## Results

### Overview of biospecimen collection and multi-omics analysis

Blood biospecimens were collected from astronauts participating in the Ax-2 and Ax-3 missions at multiple time points spanning the pre-flight and post-flight phases (Figure 1). Peripheral Blood Mononuclear Cells (PBMCs), RNA, and plasma were isolated from blood collected in different tube types. For Ax-2, samples were collected at six time points, ranging from L-90 (90 days pre-launch) to R+14 (14 days post-return), while Ax-3 included five collection points, from L-30 (30 days pre-launch) to R+60/61 (60/61 days post-return). These samples underwent comprehensive multi-omics profiling using single-cell transcriptomics (10x Genomics), bulk transcriptomics (Total RNA-seq), and proteomics (Olink Explore HT). Analytical approaches such as differential expression analysis, heatmap visualization, and principal coordinate analysis (PCA) were applied across datasets (Figure 1).

**Figure 1:**
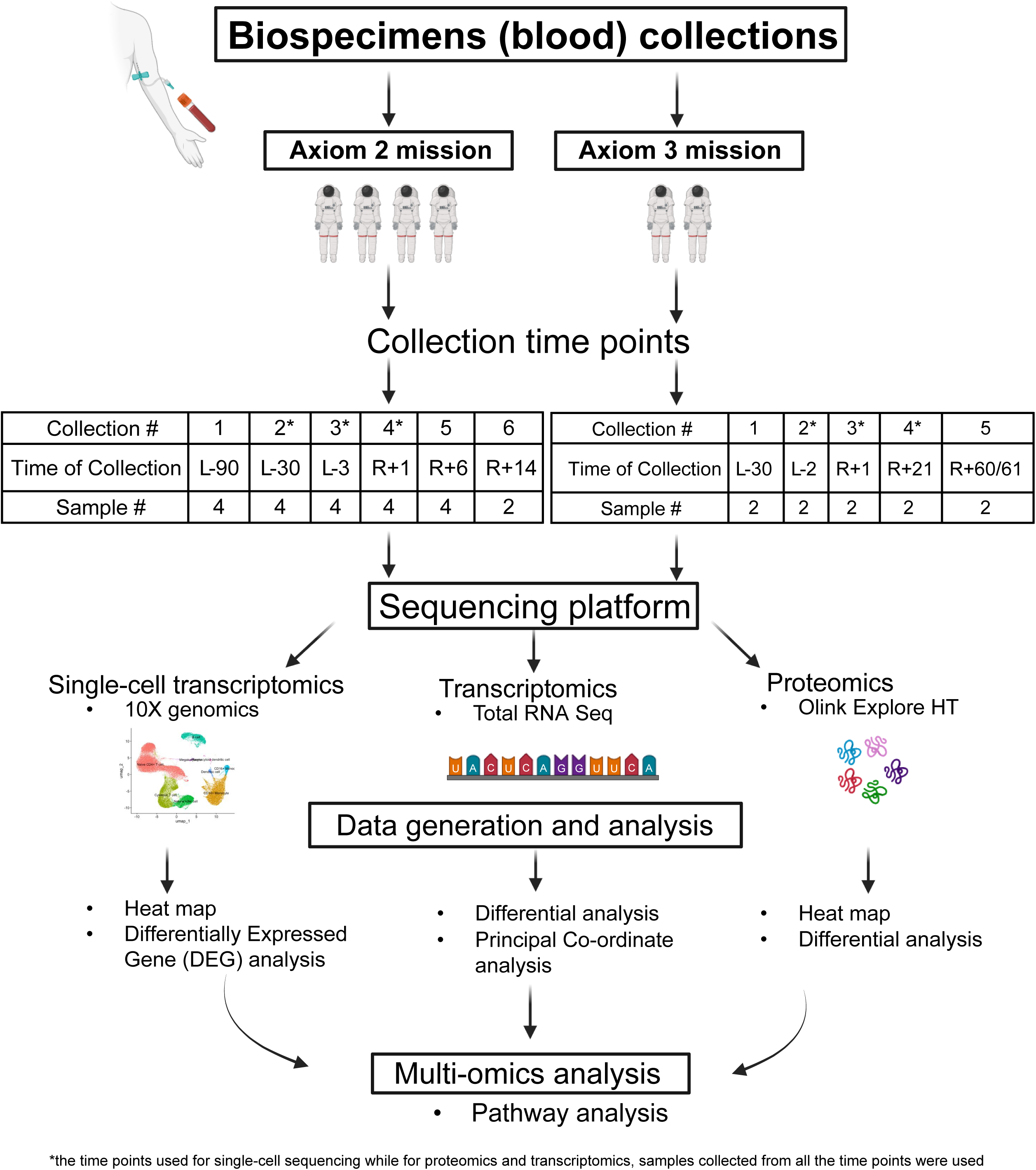
Workflow of biospecimen collection and multi-omics analysis for Ax-2 and Ax-3 missions. Blood samples were collected at multiple timepoints across both missions. Samples were processed using single-cell transcriptomics (10x Genomics), bulk transcriptomics (Total RNA-seq), and proteomics (Olink Explore HT). Data were analyzed using differential expression, heatmaps, and principal coordinate analysis. The resulting datasets were integrated through multi-omics pathway analysis to identify molecular changes associated with spaceflight. This figure was created with BioRender.com under the Baylor College of Medicine Institutional license. Time points used for single-cell sequencing*. Proteomics and transcriptomics, samples collected from all-time points.

### Single-Cell Transcriptomic Profiling Across Ax-2 and Ax-3 Spaceflight Missions

Single-cell RNA sequencing was performed on 11 samples from Ax-2, collected from four individuals at three sessions: L-30 (4 samples), L-3 (4 samples), and R+1 (4 samples), representing pre-flight, near-launch, and return phases (Figure 2A). One sample failed quality control at L-3 and therefore, only three samples were used from this time point. For Ax-3, six samples were collected from two individuals at L-2 (2 samples), R+1 (2 samples), and R+21 (2 samples), corresponding to pre-flight, early post-flight, and late post-flight phases (Figure 2B). This longitudinal and overlapping sampling design enabled the assessment of transcriptomic changes associated with spaceflight across different mission timelines. All samples from Ax-3 passed quality control thresholds and were included in downstream single-cell analysis. The single-cell QC metrics are in the supplementary Table S1.

**Figure 2:**
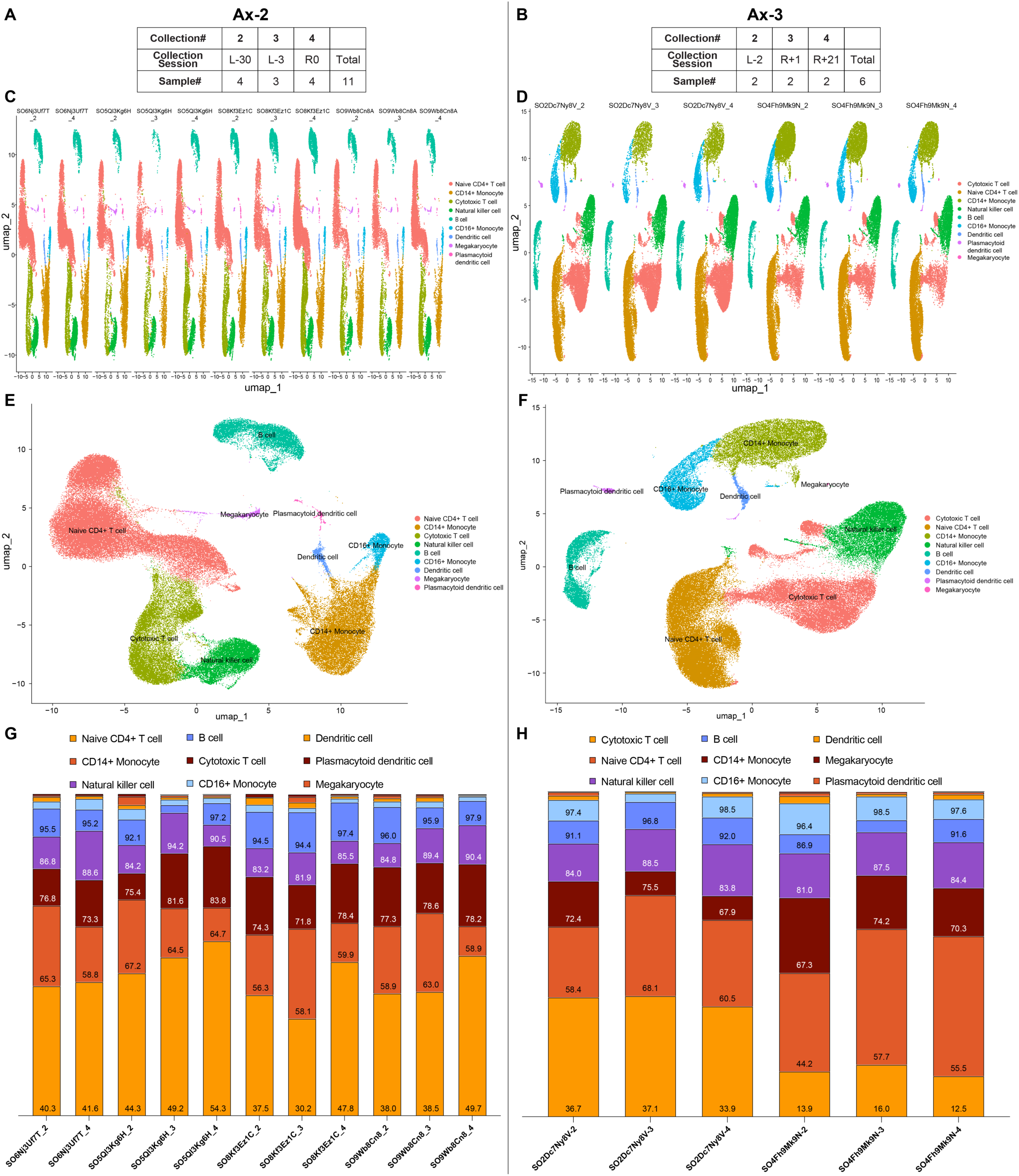
Overview of sample collection and immune cell profiling of single-cell transcriptomics data from Ax-2 and Ax-3 missions. (A, B) Summary of sample collection timepoints and total samples for Ax-2 and Ax-3. (C, D) UMAP plots illustrating clustering of immune cell types based on transcriptional profiles from Ax-2 (C) and Ax-3 (D) across individual samples. (E, F) Integrated UMAP plots from 11 samples across Ax-2 (E) and 6 samples across Ax-3 (F) for downstream functional analysis. (G, H) Bar graphs showing the distribution of immune cell types across individual samples for Ax-2 (G) and Ax-3 (H).

Dimensionality reduction using Uniform Manifold Approximation and Projection (UMAP) revealed distinct clustering patterns across samples (11 from Ax-2 and 6 from Ax-3) (Figure 2C and 2D). UMAP analysis identified 14 transcriptionally distinct clusters (0–13), including Cytotoxic T cells, Naive CD4⁺ T cells, CD14⁺ and CD16⁺ Monocytes, Natural Killer cells, B cells, Dendritic cells, Plasmacytoid Dendritic cells, and Megakaryocytes (Figure 2E and 2F). The consistent presence and separation of these cell populations across samples suggest robust cell-type identification and reproducibility of the clustering approach (Figure 2C, 2D, 2E, and 2F). Clusters were annotated using marker gene expression (Figure S1) and are the basis for analyzing spaceflight-associated biological changes. Each cell type formed well-separated clusters, and all the major cell types in Peripheral Blood Mononuclear Cells (PBMCs) were identified in all sequenced libraries. (Figure 2E and 2F).

### Longitudinal Single-Cell RNA Sequencing Uncovers Immune Cell Changes in Spaceflight

Bar plots showing the distribution of immune cell types across individual samples from the Ax-2 and Ax-3 missions revealed consistent representation of major cell populations (Figure 2G and 2H). Analysis of immune cell composition across samples from the Ax-2 and Ax-3 missions revealed distinct shifts in cell-type abundance from pre-flight to post-flight timepoints. Specifically, the proportions of CD14+ Monocytes, CD16+ Monocytes, Plasmacytoid Dendritic Cells, Dendritic Cells, and Megakaryocytes decreased following spaceflight (Figure 2G and 2H). In contrast, Naive CD4+ T cells and Cytotoxic T cells showed increased representation in the post-flight sample compared to the pre-flight sample (Figure 2G and 2H). The expression patterns of HLAs corresponding to MHC class I (HLA-A, HLA-C) and MHC class II (HLA-DRA, HLA-DRB1, HLA-DRB5), along with Beta-2 microglobulin (B2M), a light chain protein essential for the structural and functional integrity of MHC class I molecules, were evaluated. The MHC-I–associated transcripts were increased after spaceflight, whereas MHC-II–associated transcripts were decreased (Figure S2A and S2B). Notably, evaluation of individual expression profiles exhibits some variance reflecting personalized immune responses, particularly in the magnitude of MHC-I upregulation. Overall, the expression pattern remains consistent across all subjects (Figure S2C and S2D). Taken together, these shifts in immune cell composition and MHC expression highlight a coordinated remodeling of the immune landscape during spaceflight, setting the stage for transcriptome-wide analyses to identify differentially expressed genes (DEGs) and pathway-level changes underlying these cellular adaptations.

### Longer Duration Mission is associated with more differentially expressed genes (DEGs)

Transcriptomic profiling of peripheral blood mononuclear cells (PBMCs) from the Ax-2 and Ax-3 missions revealed distinct and overlapping gene expression changes in response to spaceflight. Venn diagrams showed unique and shared transcripts that were significantly up- or downregulated, highlighting both mission-specific and common transcriptional responses in Ax-2 and Ax-3 (Figure 3A, Table S2-S6).

**Figure 3:**
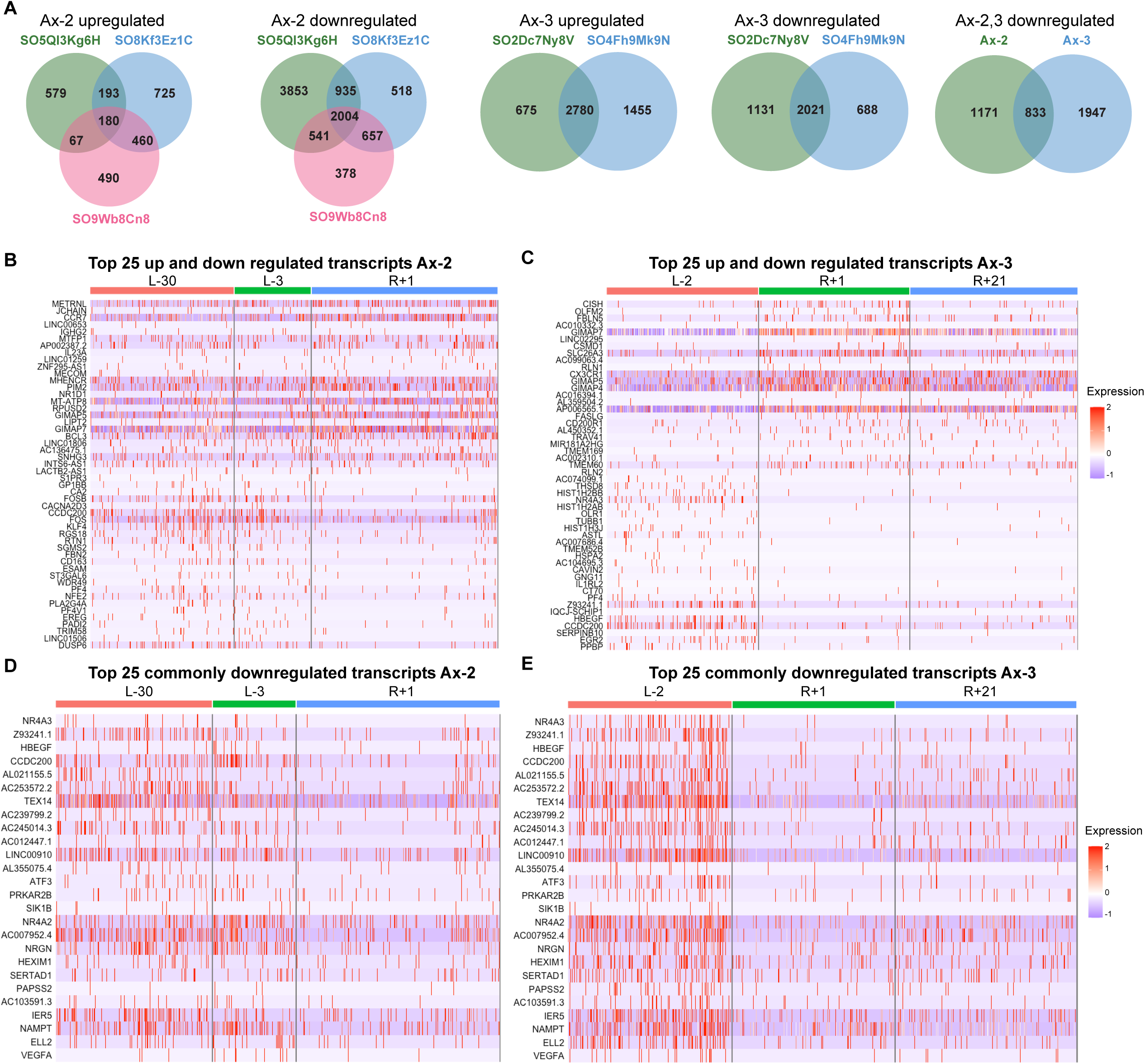
Transcriptomic analysis of PBMCs from Ax-2 and Ax-3 missions. (A) Venn diagrams showing overlap of upregulated (left) and downregulated (right) transcripts in Ax-2 and Ax-3. (B) Heatmaps of the top 25 upregulated (left) and top 25 downregulated (right) transcripts in Ax-2. (C) Heatmaps of the top 25 upregulated (left) and top 25 downregulated (right) transcripts in Ax-3. (D, E) Combined heatmap of the top 25 commonly downregulated transcripts across Ax-2 and Ax-3.

Heatmaps of the top 25 upregulated and downregulated genes for each mission showed clear separation between pre-flight and post-flight samples, indicating robust transcriptional shifts induced by spaceflight, with expression patterns beginning to normalize to pre-flight levels by the R+21 timepoint in some transcripts (Figure 3B and 3C). This gene set in Ax-2 reflects a strong acute stress–response signature, characterized by upregulation of immediate-early genes such as FOS, FOSB, KLF4, DUSP6, and NR1D1 (Figure 3B and 3C). It also includes key regulators for immune cell activation and trafficking (CCR7, GIMAP5/7, IL23A, BCL3) and markers of monocyte/macrophage and lymphocyte function. Several platelet and megakaryocyte-associated genes (PF4, PF4V1, GP1BB, NFE2, RGS18) point to hematopoietic and coagulation-related changes. This gene set in Ax-3 reflects coordinated responses to spaceflight-induced stress, encompassing extracellular matrix and vascular remodeling (SPARC, VEGFA, HBEGF), acute stress and DNA damage response (ATF3, H2AFX), immune and hematopoietic modulation (PF4, TUBB1, IL3RA, FLT3, GNG11), and metabolic adaptation (NAMPT, CA2, GMPR).

The total number of DEGs commonly upregulated in Ax-2 (180) was markedly lower than in Ax-3 (2,780). Although 2,780 DEGs were upregulated and 2,031 downregulated at R+1, these numbers dropped to 1,186 and 1,919, respectively, at R+21 in Ax-3. Notably, 833 DEGs were commonly downregulated, and 59 DEGs were commonly upregulated across both missions at the R+1 timepoint (Figure 3A). A combined heatmap of the top 25 of those 833 commonly downregulated transcripts further emphasized shared molecular signatures across missions in response to spaceflight. At the same time, the reduced magnitude of downregulation at R+21 suggests that these effects are largely reversible following return to Earth (Figure 3D and 3E).

Pathway-specific transcriptomic analysis revealed transient alterations in immune signaling pathways following spaceflight in both Ax-2 and Ax-3 missions. Dot plots representing gene expression changes across five pathways of Platelet activation, Fcγ-mediated phagocytosis, Chemokine signaling, B-cell signaling, and NF-κB signaling are shown in Figure S3A. In both cohorts, the most pronounced changes occurred at the early post-flight timepoint (R+1), with decreased expression of pathway-associated genes. By R+21, expression levels largely returned to baseline, indicating a rebound effect. Consistent with the overall group-level patterns, the changes across the five immune-related pathways were also evident at the sample level (Figure S3B). Each individual within the Ax-2 and Ax-3 groups exhibited a comparable directional trend in pathway activation, with some variation in expression across samples.

### Sequencing Metrics and Principal Component Analysis Reveal Individually Unique Transcriptomic Signatures Across Spaceflight Missions

RNA-seq (bulk) libraries for 22 Ax-2 and 10 Ax-3 samples, collected across different timepoints (Figure 1), were prepared using Illumina TruSeq Stranded Total RNA LP Globin kit and sequenced on Illumina Novaseq 6000. On average, each library yielded ∼140.3 million reads with a mean mapping rate of 92.9%. Of mapped reads, 43.9% were uniquely aligned. Most reads (94.9%) were mapped to intragenic regions with 70% aligning to exonic regions, 24.9% to intronic regions, and 5.02% of reads mapped to intergenic regions. Across all samples, on average 29,196 genes and 150,787 transcripts were detected, with a strand specificity of 96.1%. Detailed metrics for individual libraries are provided in Table S7.

Principal component analysis (PCA) of RNA-seq (bulk) data from Ax-2 and Ax-3 was used to visualize gene expression patterns across individuals and timepoints; in Ax-2, samples clustered by individual, with PC1 explaining 32%, PC2 12% and PC3 9.7% of the variance (Figure S4A). In Ax-3, PCA showed even stronger separation, with PC1 explaining 32%, PC2 16%, and PC3 14% of the variance (Figure S4B). In both the missions and across six subjects, well-defined and non-overlapping clusters were evident, highlighting distinct transcriptomic signatures between individuals.

### High-Throughput Proteomic Analysis Using Olink Explore HT Reveals PCA Visualization and Heatmap Profiling of Protein Expression Patterns

Analysis of the proteomics dataset demonstrated separation of protein-level profiles and strong clustering of individual subjects. In the Ax–2 dataset, PC1, PC2, and PC3 captured 22.7%, 12.5%, and 11.7% of the variance, respectively (Figure S4C). For Ax-3 analysis, it accounted for 29.9%, 15.4%, and 10.4% of the variance, with similar clustering of individual subjects (Figure S4D).

To assess longitudinal changes in protein expression, high-throughput proteomic profiling was performed using the Olink Explore HT platform, which quantifies approximately 5,400 proteins per sample. Global heatmaps were generated to provide an overview of the proteomic landscape, enabling visualization of temporal patterns and shifts in protein abundance. A total of 32 samples were collected from two missions, 22 samples from Ax-2 and 10 samples from Ax-3 (Figure 4E and 4F). Each sample was analyzed using longitudinal sampling across multiple timepoints for individual participants. Statistical analysis identified a substantial number of significantly altered proteins across timepoints: In the Ax-2 mission samples, 1,112 proteins were significantly different across timepoints and in the Ax-3 mission samples, 559 proteins showed significant changes. The heatmap for Ax-2 (Figure S4E and S4G) shows proteomic profiles of 22 samples collected at six timepoints: L-90, L-30, L-3, R+1, R+6, and R+14 whereas for Ax-3 (Figure S4F and S4H), a total of 10 samples were collected at five different timepoints: L-30, L-2, R+1, R+21, and R+60/61. Interestingly, for the Ax-2 mission, the L-3 timepoint, which was three days before launch, showed the most significant proteomic changes. The initial hypothesis was that this could be due to either a stress response from the impending launch or a discrepancy in sample handling or collection. However, when compared to the Ax-3 datasets, this pattern did not align with either explanation. Due to this uncertainty, the L-3 proteomics data from the Ax-2 mission was excluded from the timepoint analysis. One of the key observations from both missions is that protein expression levels tend to be elevated immediately after returning from space, compared to both pre-flight and later post-flight timepoints. This trend highlights a transient but pronounced biological response to spaceflight re-entry.

**Figure 4.**
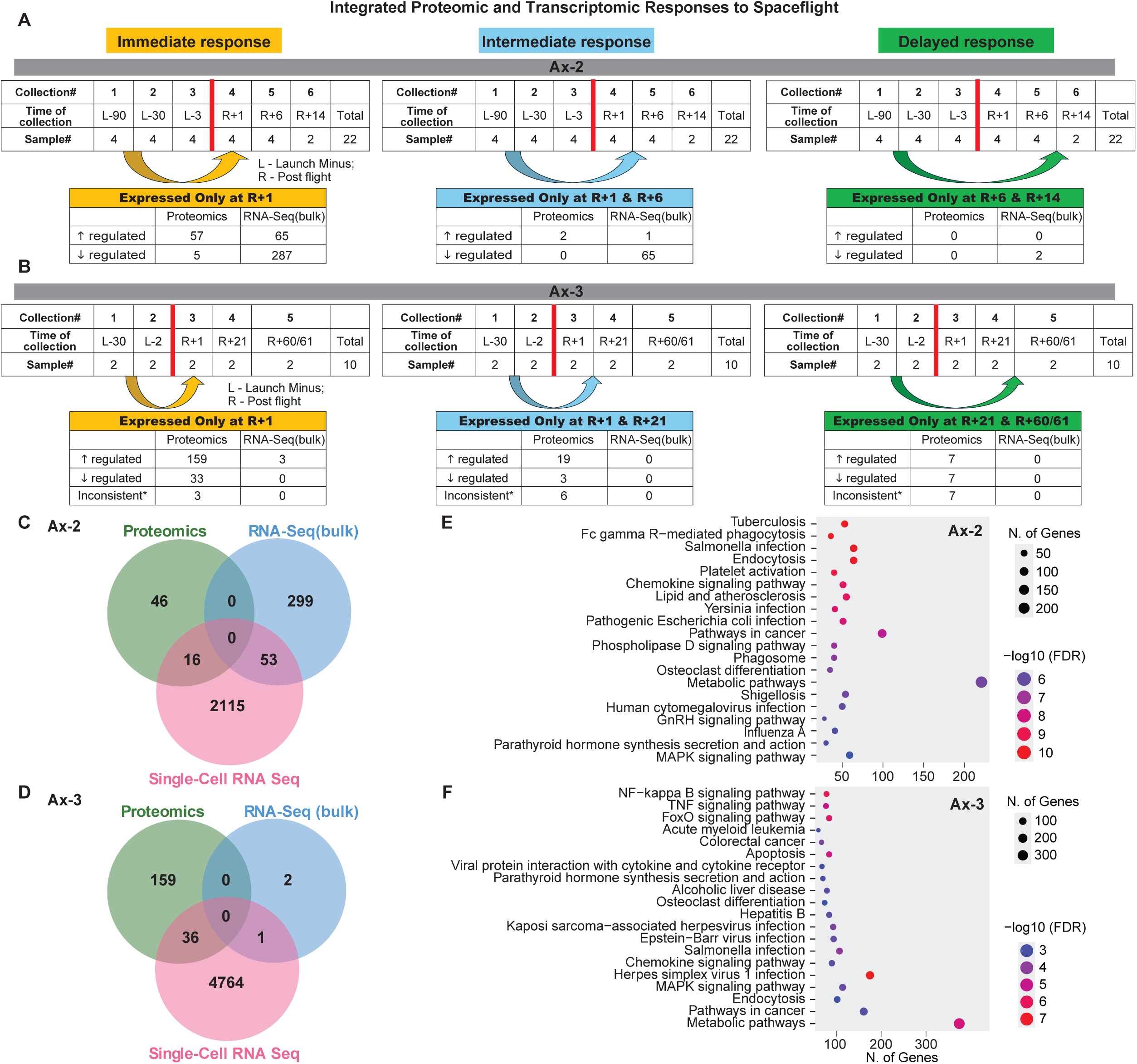
Integrated multi-omics responses to spaceflight. (A, B) Proteomic and transcriptomic profiles from Ax-2 and Ax-3 reveal distinct molecular phases: Immediate (R+1), Intermediate (Ax-2: R+1/R+6; Ax-3: R+1/R+21), and Delayed (Ax-2: R+6/R+14; Ax-3: R+21/R+60/61). The red vertical line in the table indicates a separation between pre and post-flight.(C, D) Venn diagrams showing overlap among proteomics, RNA-seq (bulk), and single-cell RNA-seq captured the most unique features, highlighting the complementarity of platforms. (E, F) Top 20 spaceflight responsive pathways across Ax-2 and Ax-3 from proteomics, RNA-seq (bulk), and single-cell RNA-seq; dot size indicates gene count, color reflects significance.

### Integrated analysis of proteomic and RNA-Seq (bulk) data from two spaceflight missions reveals distinct molecular response phases

To investigate the biological effects of spaceflight, high-throughput proteomic and RNA-seq (bulk) data were analyzed from two independent missions, Ax-2, and Ax-3. The analysis focused on identifying molecular responses by comparing post-flight samples to pre-flight baselines. Since data from both missions were collected and generated at the same timepoints, it enabled a simultaneous comparison of changes in protein and gene expression over time. These responses were categorized into three categories: Immediate, Intermediate, and Delayed responses based on the timepoints at which specific proteins or transcripts were uniquely expressed. Immediate response refers to proteins and transcripts that show differential regulation at timepoint R+1 compared to L-90 and L-30. Intermediate response is based on changes observed at R+1 and R+6, relative to L-90 and L-30. Delayed response involves changes detected at R+6 and R+14, also compared to L-90 and L-30 (Figure 4A and B).

In Ax-2, the immediate response group includes 57 upregulated and 5 downregulated proteins, along with 65 upregulated and 287 downregulated transcripts (Figure 4A and Table S8). Proteomics and RNA-Seq (bulk) analysis of Ax-2 samples identified upregulation of proteins FOSB (log2FC: 4.5), TSC22D3 (2.5), CDC42BPB (2.6), KIFC3 (2.6), SNX29 (2.9), PRKG1 (3.1) and transcripts COL4A1 (2.4), and ATP1A2 (2.5). Downregulated proteins include ITM2A (−1.6), TMPRSS15 (−1.1), CLEC6A (−0.7), TNFSF11 (−0.7), and GAL (−0.5) alongside transcript decrease in HBQ1 (−2.0), ALAS2 (−1.9), and IGHA1 (−1.7) (Table S8). The intermediate phase showed only two upregulated proteins and one upregulated transcript, alongside 65 downregulated transcripts. The delayed response revealed no uniquely expressed proteins but included statistically significant downregulation of two transcripts (p < 0.05). These findings show that significant changes are seen immediately after returning from space (after splash down) and become transient over time (Figure 4A and Table S8).

In Ax-3, the immediate response group (R+1) includes 159 upregulated and 33 downregulated proteins, and three upregulated transcripts. Three proteins showed inconsistent expression (Figure 4B and Table S9). Inconsistent proteins and transcripts were defined as those displaying divergent expression patterns across timepoints, such as upregulation at one timepoint and downregulation at another. Proteomic profiling of the top 25 proteins at R+1 showed differential expression compared to pre-flight baselines (L-30 and L-2). Proteomics and RNA-Seq (bulk) analysis of Ax-3 samples showed upregulation of proteins including FGF21 (log2FC: 2.8), PTTG1 (5.4), RNF31 (6.0), DOC2B (6.3), ITGAX (6.5), and MAGEA3/MAGEA6 (6.7) and a few upregulated transcripts, which include S100A12 (1.7), ANXA3 (1.4) and LMNB1 (1.2). A subset of proteins and transcripts plays key roles in cellular senescence and may have significant implications for Spaceflight. Downregulated proteins include CTAG1A/CTAG1B (log2FC: −5.6), FBXL5 (−4.8), and ATP5PO (−2.3) (Table S9). The Intermediate phase, defined by differential expression at R+1 and R+21, includes 19 upregulated and 3 downregulated proteins, with six proteins showing inconsistent expression (Figure 4B and Table S9). No significant transcriptomic changes were detected in this phase. The delayed response, observed at R+21 and R+60/61, involved 7 upregulated and 7 downregulated proteins, along with 7 inconsistent proteins, while RNA-Seq (bulk) data showed no significant changes (Figure 4B and Table S9).

Analysis of non-coding RNA and mitochondrial RNA-Seq datasets from Ax-2 showed non-coding RNA expression in the immediate response, with 18 transcripts upregulated and one transcript downregulated (Table S10). At R+1 and R+6 (intermediate response), only one non-coding RNA was upregulated, while R+6 and R+14 (delayed response) showed no differential expression. In contrast, mitochondrial genes showed no differential expression across immediate, intermediate, or delayed responses. In the Ax-3 data, using the above-described criteria of finding immediate, intermediate, or delayed response patterns, neither non-coding RNAs nor mitochondrial RNAs were significantly differentially regulated.

### Integrative analysis of Proteomics (OLINK), RNA sequencing (bulk), and Single-cell transcriptomics

#### Multi-Omics Comparison Reveals Distinct and Complementary Insights

A comparative analysis of proteomics (OLINK), RNA-seq (bulk), and single-cell RNA-seq was performed on post-flight samples to assess overlap in detected proteins and genes to show shared and unique features across datasets. In Ax-2, proteomics identified 46 differentially expressed proteins (DEPs), RNA-seq (bulk) detected 299 genes, and single-cell RNA-seq revealed 2,115 genes (Figure 4C and Table S11).

We observed 16 overlapping protein–transcript pairs between proteomics and single-cell RNA-seq, and 53 overlapping transcripts between bulk RNA-seq and single-cell RNA-seq. Approximately 90% of the transcripts were concordant between bulk RNA-seq and single-cell RNA-seq, whereas only ∼6% of the protein–transcript pairs showed agreement. Concordance here refers to up or down regulation in both assays. In the Ax-3 dataset, proteomics identified 159 DEPs, RNA-seq (bulk) only 2 DEGs, and single-cell RNA-seq 4,764 (Figure 4D and Table S12). There were 36 overlapping DEPs/DEGs between proteomics and single-cell RNA-seq, and concordance was approximately 50%. Additionally, one overlapping RNA-seq (bulk) and single-cell RNA-seq was discordant. There were no common overlapping proteins/transcripts across all three platforms in both missions.

### Multi-Omics Analysis Reveals Immune and Metabolic Pathways Consistently Impacted Across Spaceflight Missions

Pathway enrichment analysis was conducted using ShinyGo 0.80 to identify biological processes significantly impacted by spaceflight. The top 20 pathways for each dataset for Ax-2 and Ax-3 mission were ranked by the number of genes involved and statistical significance (−log10 FDR) (Figure 4E and F).

In Ax-2, pathways such as tuberculosis, Fc gamma R-mediated phagocytosis, Salmonella infection, and platelet activation were among the most enriched, with gene counts ranging from 50 to 200 (Table S13). These pathways suggest activation of immune-related and inflammatory processes. Additional pathways, including lipid and atherosclerosis, osteoclast differentiation, phagosome, and metabolic pathways, indicate alterations in lipid metabolism and cellular homeostasis. In Ax-3, enrichment was observed in pathways such as NF-kappa B signaling, acute myeloid leukemia, TNF signaling, and apoptosis, with some pathways involving over 300 genes (Table S14). The top 20 common pathways across missions include endocytosis, MAPK signaling, Salmonella infection, which share many genes and are involved in immune and stress responses, osteoclast differentiation, and metabolic pathways (Figure 4E and F).

To examine spaceflight-related biological effects, particularly those linked to NASA HRP priority areas (HRP-47052) and known phenotypic changes, top pathways with broad impact were further analyzed: (1) Osteoclast differentiation, (2) NF-κB signaling, and (3) Coagulation, Complement, and (4) Platelet Activation.

### Osteoclast Differentiation Signaling Pathway

Differentially expressed proteins in the osteoclast pathway were detected across Ax-2 and Ax-3 multi-omics datasets, including key regulators such as TRAF2, FOSB, and RANKL (TNFSF11), along with co-stimulatory molecules c-Fos, MITF, Gab2, Syk, and PLCγ (Figure 5A and Table S15). In Ax-2 proteomics data, FOSB showed an increase (∼4.5 fold) and TRAF2 was elevated, indicating activation of osteoclastogenesis and enhanced RANK signaling. In contrast, RANKL was moderately reduced in Ax-2 (−0.7) and further in Ax-3 (−1.8) proteomics. Single-cell datasets revealed consistent declines in several regulators: c-Fos (−1.9 Ax-2; −0.9 Ax-3), MITF (−1.6 Ax-2; −0.9 Ax-3), Gab2 (−1.0 Ax-2; −1.3 Ax-3), Syk (−0.7 Ax-2; −0.4 Ax-3), and PLCG2 (−0.6 Ax-2; −0.4) Together, these results suggest a disruption of the balance between osteoclast and osteoblast activity.

**Figure 5.**
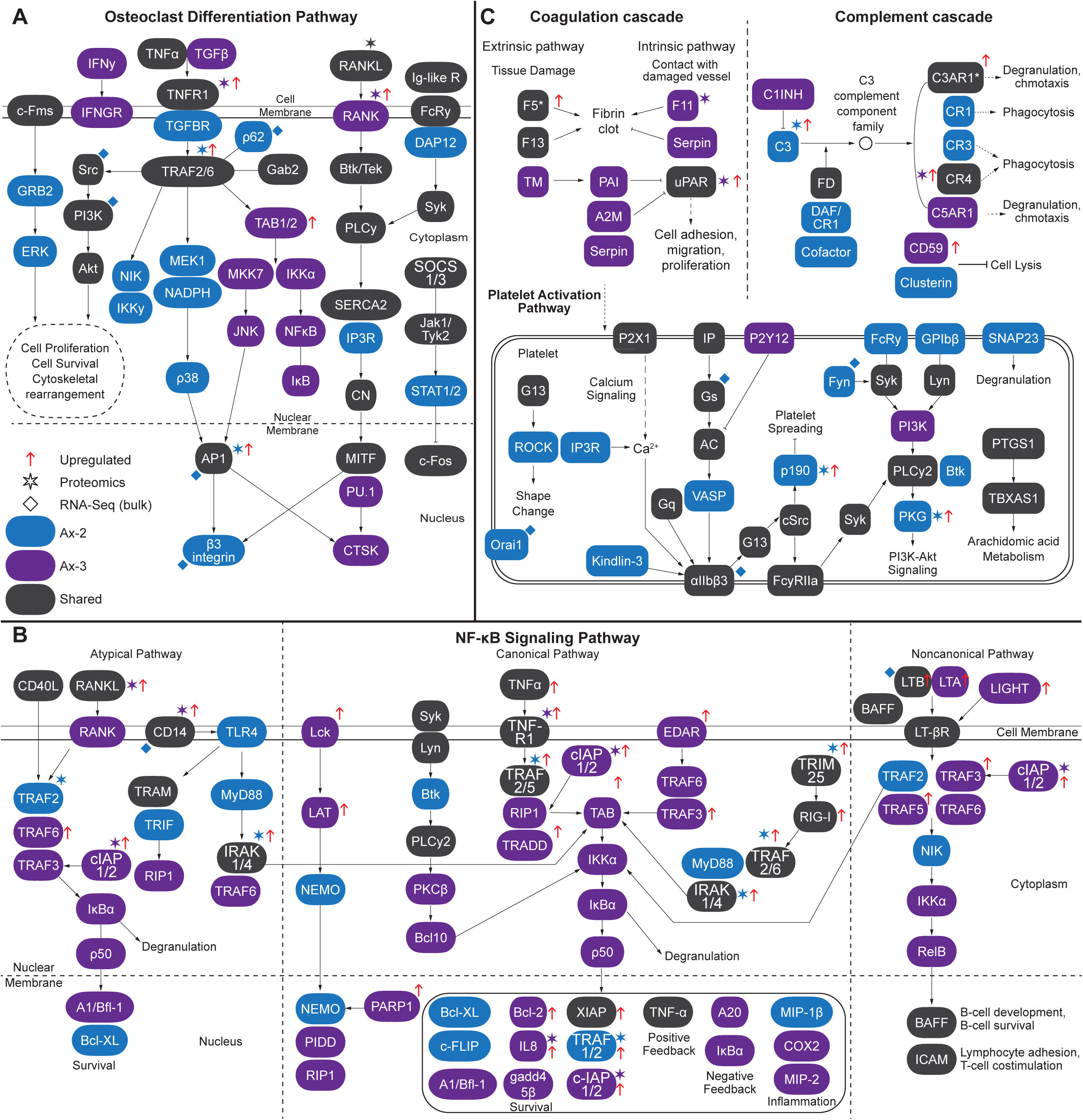
Spaceflight-induced modulation of key signaling pathways. (A) Osteoclast differentiation, (B) NF-κB signaling, and (C) Coagulation, Complement, and Platelet Activation Pathways. Genes are color-coded by mission (Ax-2: blue; Ax-3: purple; shared: black). Most genes were identified via single-cell RNA-seq; stars denote proteomics data; diamonds denote RNA-seq (bulk); upwards pointed arrow denotes upregulation, while no arrow sign denotes downregulation. This figure was hand-curated in Adobe Illustrator 2025 using the KEGG pathway backbone to highlight the pathway-level responses to space flight.

### NF-κB Signaling Pathway Modulation in Spaceflight

Differentially expressed NF-κB signaling pathway components were detected across Ax-2 and Ax-3 multi-omics datasets (Figure 5B and Table S16). Shared regulators that showed an increase were CD40LG (Ax-2 +0.5; Ax-3 +0.9) and decreased in TNF-family ligands such as RANKL/TNFSF11 (Ax-2 −0.7; Ax-3 −1.8), LTBR (Ax-2 −1.1; Ax-3 −0.6), and TICAM2 (Ax-2 − 1.0; Ax-3 −0.4). Intracellular adaptors and kinases required for NF-κB activation, including Lyn (Ax-2 −0.8; Ax-3 −1.1), Syk (Ax-2 −0.7; Ax-3 −0.4), and PLCG2 (Ax-2 −0.6; Ax-3 −0.4), were consistently reduced, whereas IRAK1 decreased in single-cell (−0.5) but increased in proteomics (1.1). In Ax-3, single-cell data revealed downregulation of core NF-κB components IκBα (−0.6), RelB (−1.1), and downstream targets A20/TNFAIP3 (−1.3), COX 2 (−1.9), GADD45B (−2.2), and CXCL2 (−1.7), while proteomics revealed increased IL 8 (0.9) despite its reduction in single-cell (−1.6). Conversely, several adaptors and survival factors increased in TNFRSF11A (+0.7), TRAF3IP3 (+0.6), BIRC2 (+1.2), TRADD (+0.7), Lck (+0.6), and TNF-family ligands LTA (+0.7) and LIGHT/TNFSF14 (+1.0). Ax-2 mission specific changes reflected upstream dampening of receptor-level signals, including TLR4 (−1.3), TICAM1 (−0.4), and MYD88 (−0.6), along with reductions in adaptors Bcl-XL (−0.7), Btk (−0.7), NEMO (−0.7), and c-FLIP (−0.4). Proteomics revealed a notable increase in TRAF2 (+1.4). Overall, Ax-2 exhibited a weaker and more localized NF-κB response compared to Ax-3, and NF-κB signaling did not rank among the top 20 pathways in Ax-2.

### Molecular Signatures of Coagulation, Complement, and Platelet Activation pathways

Spaceflight broadly altered coagulation and hemostasis pathways, with evidence from both proteomics and single-cell datasets (Figure 5C and Table S17). Shared regulators that were downregulated included CFD (Ax-2 −0.9; Ax-3 −0.7) and F13A1 (Ax-2 −1.7; Ax-3 −0.9) in single-cell data. PLAUR/uPAR (−1.2) and ITGAX (−1.0) decreased in Ax-2 single-cell profiles (−1.2) but increased in Ax-3 proteomics for PLAUR/uPAR (0.5) and ITGAX (6.5). Ax-2 proteomics showed elevated C3 (0.8), while single-cell data revealed declines in CR1 (−0.6) and ITGAM/CR3 (−1.0). Ax-3 shows broad suppression of anticoagulant and fibrinolytic mediators (THBD −1.4; SERPINB2 −2.4) alongside selective upregulation of immune regulators (CD59 +0.6), suggesting a shift toward pro-coagulant and immune-altered states during spaceflight (Figure 5C and Table S17).

Platelet activation pathways were similarly suppressed, with single-cell data showing consistent downregulation of P2RX1 (Ax-2 −0.9; Ax-3 −1.0), PTGIR (Ax-2 −1.2; Ax-3 −0.4), SRC (Ax-2 −1.3; Ax-3 −1.1), and PLCG2 (Ax-2 −0.6; Ax-3 −0.4). In the Ax-2 proteomics data, inhibitory regulators, ARHGAP35 increased by 2-fold and PRKG1 by 3.1, suggesting active suppression of platelet activation, whereas Ax-3 showed a reduction in signaling components like PIK3R1 (−0.7) and P2RY12 (−2.2) in single-cell datasets (Figure 5C and Table S17). Collectively, these findings indicate that spaceflight disrupts multiple components of hemostasis, reducing platelet activation potential and altering complement-coagulation balance, with mission-specific regulatory differences.

## Discussion

The present study provides the first integrated multi-omics view of early human molecular responses to spaceflight across two commercial missions, Ax-2 and Ax-3, utilizing synchronized biospecimen collection outlined in the GENESTAR protocol^6^. By jointly analyzing single-cell transcriptomics, bulk RNA-seq, and high-throughput proteomics, we reveal a coordinated set of immune, metabolic, and vascular adaptations that arise within the first three weeks of spaceflight. Despite differences in mission duration and individual-level variability, several conserved signatures emerge across platforms, highlighting reproducible early-phase spaceflight responses.

The single-cell RNA-Seq profiling revealed distinct gene expression changes between pre-flight and post-flight samples (Figure 3A). The number of differentially expressed genes (DEGs) was notably higher in the longer-duration Ax-3 mission, suggesting that mission length may influence the magnitude of transcriptomic response (Figure 3A). Heatmap and pathway-specific analyses highlighted transient downregulation of immune signaling pathways such as NF-κB, B-cell signaling, and chemokine signaling immediately after returning, with partial recovery by later post-flight timepoints (Figure S3A and S3B). These patterns reflect an acute but reversible modulation of immune function following spaceflight. RNA-seq (bulk) and proteomics analysis further supported these findings. These observations relate to previous studies showing decreased metabolic activity and mitochondrial stress in PBMCs during ISS missions^10,11^.

Longitudinal analysis demonstrated post-flight shifts in immune composition, with increased representation of T cell subsets and decreased abundance of monocytes and dendritic cells (Figure 2G and 2H; Figure S1). These findings align with prior reports from the NASA Twins Study and other spaceflight investigations, which describe immune dysregulation and T-cell functional impairment under microgravity, including altered activation and cytokine signaling^4,10,12–14^. Analysis of MHC gene expression in our data further validated these shifts: MHC-I molecules (HLA-A, HLA-C, B2M), expressed on all nucleated cells, were consistently upregulated post-flight, while MHC-II molecules (HLA-DRA, HLA-DRB1, HLA-DRB5), primarily found on professional antigen-presenting cells, were markedly downregulated (Figure S2), reflecting reduced antigen presentation capacity^15^. Collectively, the observed coordinated immune shifts highlight a transient suppression of innate immune components (CD14+ and CD16+ monocytes, plasmacytoid and conventional dendritic cells) and redistribution of antigen-presentation machinery post-flight, paired with adaptive mobilization of cytotoxic and naive T cells. These findings are consistent across Ax-2 and Ax-3 and may represent universal early responses to spaceflight exposure.

Integrated proteomic and transcriptomic profiling across Ax-2 and Ax-3 reveals rapid yet transient molecular responses to short-duration spaceflight. Both missions showed pronounced immediate changes in post-flight biospecimens, dominated by stress, immune activation, and cytoskeletal remodeling, while intermediate and delayed phases exhibited minimal alterations, indicating recovery toward baseline (Table S8 and S9). These dynamics suggest that early adaptations, such as mitochondrial stress signaling and inflammatory pathways, are critical for adjusting to spaceflight. Understanding these temporal changes is essential for designing countermeasures for the long-term duration of spaceflights by targeting such early responders.

In the Ax-2 proteomics data, TSC22D3 (GILZ) (log2FC: 2.53) and FOSB (4.50) are upregulated. Microgravity can transiently induce AP-1 factors such as FOSB, while concurrently elevating TSC22D3 (GILZ) as part of a compensatory glucocorticoid-linked anti-inflammatory response^16,17^. Upregulation of COL4A1 transcripts in the RNA-Seq (bulk) data (log2FC: 2.42), which encodes a key type IV collagen, reflects basement-membrane and ECM remodeling, and previously, another gene from this family, COL4A4, was reported as differentially regulated in long-term exposure to space microgravity^18^. Elevated ATP1A2 gene expression (log2FC: 2.5) likely reflects altered Na⁺/K⁺-ATPase–mediated ion transport under microgravity, previously reported ^19^, relating to altered ion transport. The downregulation of proteins CTAG1A/CTAG1B (log2FC: −5.65), FBXL5 (−4.8), and ATP5PO (−2.3) was observed in Ax-3 (Table S9) but has not been previously reported in astronaut or rodent datasets. Functionally, CTAG1A/CTAG1B is related to immunity^20,21^. FBXL5 is an iron/oxygen-sensing E3 ligase^22,23^. Decreased ATP5PO (−2.3) in Ax-3, which encodes a subunit of ATP synthase, therefore agrees with suppression of mitochondrial OXPHOS and ATP-synthase components during spaceflight^24,25^. This also agrees with the OXPHOS suppression reported in the Inspiration4 and Twins study^4,26^. But our data conclusively provides such molecular details.

RNA-seq (bulk) from the Ax-3 mission showed significant upregulation of LMNB1, a core senescence marker^27,28^, whose overexpression can induce senescence^29^. Additionally, the proteomics analysis indicated upregulation of proteins involved in regulating cellular senescence, including FGF21, PTTG1, and DOC2B, with elevated DOC2B previously linked to cell-cycle arrest and senescence^30^ (Table S9). Because unregulated senescence can transition from tumor suppressive to pro-tumorigenic and pro-aging^31^, these changes suggest that Space radiation and microgravity may accelerate organismal senescence. Ongoing development of senolytic therapies may help mitigate senescent cell accumulation during long-duration missions^32–35^. Collectively, these observations suggest that short-duration spaceflight can trigger stress-response and senescence-related pathways, with greater molecular perturbation associated with increased mission duration, as observed in Ax-3.

Several of the upregulated Proteins listed in Table S9 such as ITGAX (6.49), RNF31 (log2FC: 6.02), MAGEA3/6 (6.7), DOC2B (6.0), PTTG1 (5.39)^21,24^ FGF21 (2.8) (Table S9) are potential biomarkers as they are part of the pathways that govern the mitochondrial and lipid stress reported in rodent spaceflight liver study and reflect mitochondrial and lipid stress^36^. DOC2B (6.26) participates in secretion pathways, though it is new to spaceflight^37^ (Table S9).

Several long non-coding RNA (lncRNA) transcripts identified from RNA-Seq (bulk) data, including LINC00461 (2.5), HAND2-AS1 (2.5), and HULC (1.6), play a critical role in gene regulation and stress adaptation (Table S10). Recent studies report that exosomal lncRNAs are significantly altered in astronauts after space missions, implicating pathways related to cardiovascular health, immune regulation, and cancer risk^38^. These findings, together with our observed lncRNA upregulation, suggest that non-coding RNAs may serve as biomarkers for spaceflight.

The comparative multi-omics analysis revealed limited overlap in differentially expressed proteins and genes across platforms, emphasizing the unique contributions of each method. This lack of overlap across all three assays highlights the importance of integrative approaches to fully characterize complex biological responses and a way to maximize discovery from a limited number of samples (Figure 4C and D).

Pathway enrichment patterns in Ax-2 and Ax-3 suggest that spaceflight triggers system-level changes. The prominence of immune infection pathways (e.g., Salmonella infection, Fc gamma R-mediated phagocytosis) in Ax-2 likely reflects an acute innate immune change ^21^ in addition to these pathways having several shared genes. In contrast, Ax-3 enrichment of NF-κB signaling, TNF signaling, and Apoptosis points to further immune suppression and stress-response remodeling, as this Ax-3 mission is twice as long in duration when compared to Ax-2. Shared pathways such as Osteoclast differentiation and metabolic processes underscore conserved effects (Figure 4E and F). These signatures highlight the need for countermeasures targeting immune resilience and skeletal integrity during extended missions.

Spaceflight appears to perturb bone remodeling by altering osteoclast signaling. Strong upregulation of FOSB in Ax-2 and elevated TRAF6 indicate activation of transcriptional programs and RANK signaling, key drivers of osteoclastogenesis. Conversely, downregulation of c-Fos, MITF, Gab2, Syk, and PLCy in single-cell RNA-seq and RANKL in proteomics suggests an opposite response, potentially disrupting osteoclast–osteoblast balance (Figure 5A and Table S15). While bone resorption and altered osteoclast markers have been documented in rodent spaceflight and simulated microgravity models, the contrasting transcriptional activation and signaling suppression observed here underscore a more nuanced disruption of osteoclast–osteoblast balance during the early days of human spaceflight^39,40^. We also noticed a downregulation of FOSB in single-cell RNA-seq data. The observed discordance, lower transcript levels, but higher protein abundance, suggests post-transcriptional regulation^41^.

Our data reveal a coordinated suppression of NF-κB signaling during spaceflight, most pronounced in Ax-3, which is twice as long as the Ax-2 mission (Figure 4B and Table S16). Downregulation of upstream receptors (TLR4, MYD88), TNF-family ligands (RANKL, BAFF), and core components (IκBα, RelB, IKKα) suggests broad inhibition of inflammatory and stress-response pathways. Reduced expression of kinases (Lyn, Syk, PLCG2) and downstream targets (A20/TNFAIP3, COX-2, GADD45B) further supports impaired immune activation. In contrast, Ax-2 exhibited a more limited response, affecting fewer genes and different nodes within the network. These findings align with prior reports of diminished T-cell and innate immune signaling under microgravity and simulated conditions^42^.

Spaceflight induces systemic hemostatic changes, as shown by coordinated modulation of coagulation, complement, and platelet activation pathways in Ax-2 and Ax-3. Acute plasma-proteome shifts have been documented in astronauts after short-duration spaceflight, including alterations in coagulation, oxidative stress, and neuro-associated pathways^43^. Platelet signaling genes (P2RX1, PTGIR, GNAS, SRC) were suppressed, with Ax-2 showing increased inhibitory regulators (ARHGAP35, PRKG1), while in Ax-3, it was adhesion molecules (ITGA2B, PIK3R1) (Figure 5C and Table S17). These findings align with studies reporting microgravity-associated coagulation imbalance^44^ and clinical evidence of venous thrombosis during ISS missions ^45^. This also underscores the importance of early timepoint sampling for detecting spaceflight-associated molecular changes. Because in-flight blood collection poses significant logistical challenges, recent efforts have utilized alternative approaches for biospecimen collection to enable proteomic profiling^46^.

In conclusion, this study provides the molecular explanation and new biomarkers for physiological changes associated with the first three weeks of spaceflight, while confirming previously reported physiological responses. Importantly, through an integrated multi-omics approach, we identified conserved molecular signatures across two Axiom missions, demonstrating the robustness of the standardized biospecimen collection and the generated data. This cross-mission consistency highlights the stability of these signatures and their potential biological relevance for spaceflight adaptation. The insights provided in this study lay a foundation for developing targeted countermeasures to mitigate immune, skeletal, and vascular complications associated with spaceflight.

### Limitations and cautious interpretation

- Small cohort size limits statistical power and generalizability. Findings should be interpreted as descriptive and hypothesis-generating rather than definitive.
- Pre-launch collections at KSC and operational stress (sleep disruption, travel, training load, and anticipatory stress) may contribute to peri-flight signatures; the Ax-2 L-3 proteomics anomaly underscores potential pre-analytical variability.
- The analysis focused on molecular signals and not clinical endpoints.

## Resource availability

### Data availability

Datasets from Single-cell RNA-seq, RNA-seq (bulk), and Proteomics (OLINK) have been uploaded to the TRISH EXPAND TrialX database.

### Code availability

The R script used in the OLINK analysis on this manuscript is available on GitHub^47^ (https://github.com/qiaoyanw/Olink-Explore-HT-Analysis-Methods-for-TRISH_AX2_AX3-Flight/tree/main)

## Acknowledgements

This study was funded (Grant# INN0010) by the Translational Research Institute for Space Health through NASA Cooperative Agreement NNX16AO69A. Single-Cell libraries were prepared by the Single-Cell Genomics Core at Baylor College of Medicine, which is supported with funding from the CPRIT RP200504 and P30CA125123. The authors are grateful to the study participants and production teams at the Human Genome Sequencing Center for data generation. We thank Dr. Audra Iness for the critical review of this manuscript.

## Author Contributions

Conceptualization: H.D, A.S, Z.M; Data Generation: Z.M, A.K, K.K, S.V.B, H.C, Q.J, M.C.G; Data Analysis: A.S, Z.M, X.Q, K.W, Q.W, D.K; Writing: H.D, A.S, Z.M; Review & Editing: Z.M, A.S, K.W, D.K, K.K, A.K, M.M.M, E.U, S.A.A, M.G, M.A.M.M, J.W, D.M, R.A.G and H.D.

## DECLARATION OF INTERESTS

The authors declare no conflict of interest.

## STAR★METHODS

- **KEY RESOURCE TABLE**
- **METHOD DETAILS**

## Methods

### IRB statement on human subjects research

Protocol title: EXPAND-MESH (Enhancing exploration platforms and analog definition - multimodal evaluation of spaceflight participant health. (H-52035).

### Biospecimen Collection

The biospecimens were collected at time points L-90, L-30, L-3, R + 1, R + 6 and R + 14 from Ax-2 (n=4) and at L-30, L-2, R+1, R+21 and R+60/61 from Ax-3 (n=2). All but immediate pre-launch collections, L-3 (Ax-2) and L-2 (Ax-3), happened at Axiom Space Inc. headquarters in Houston, Texas. The biospecimens were shipped within ∼2 hours of collection to the Human Genome Sequencing Center (HGSC) at Baylor College of Medicine (BCM), Houston, Texas, for initial processing. Samples from the L-3 (Ax-2) and L-2 (Ax-3) time point were collected at Kennedy Space Center (KSC), Florida, processed at the Burnett School of Biomedical Sciences (BSBS), University of Central Florida, and then shipped to BCM-HGSC for further processing and biobanking as per the GENESTAR manual^6^.

### PBMC Isolation, Sample Collection Methods

To isolate human peripheral blood mononuclear cells (PBMCs), blood samples were collected in BD cell preparation tubes as previously described in the GENESTAR manual^6^. To count the cells, 50 µL cell suspension was transferred to a new 1.5-mL tube and mixed with 50 µL trypan blue (Sigma-Aldrich Cat. No. 72-57-1) at a 1:1 ratio. 10 µL of the mixture was loaded onto the hemocytometer (Hausser Scientific). Then, the hemocytometer was placed on the stage of the Corning CytoSMART Cell Counter, optimally focused, and visualized using the Axion Biosystems software. Using the built-in feature of the software, cells were gated based on cell morphology, size, and trypan blue exclusion to determine total, live, and dead cell density, along with percent viability. Upon completion of cell counting, an equal volume of freeze media (30% DMSO, 40% FBS, and 30% RPMI) was then added to the original PBMC suspension, and cells were frozen in a Mr. Frosty container at –80°C.

### Single-cell RNA Sequencing

Single-cell 5’ Gene Expression Library was prepared according to Chromium Single Cell Immune Profiling Solution 5’ v2 and 5’ v3 (10x Genomics) for samples from Ax-2 and Ax-3, respectively. In brief, single cells, reverse transcription (RT) reagents, Gel Beads containing barcoded oligonucleotides, and oil were loaded on a Chromium X (10x Genomics) to generate single-cell GEMS (Gel Beads-In-Emulsions) where full-length cDNA was synthesized and barcoded for each single cell. Subsequently, the GEMS was broken, and cDNA from each single cell was pooled. Following cleanup using Dynabeads- MyOne Silane Beads, the cDNA was amplified by PCR. The amplified cDNA was then fragmented to an optimal size, and the 5’ Gene Expression (GEX) library was generated via End-repair, A-tailing, adaptor ligation, and PCR amplification. Libraries were quantified using tape station and qPCR, pooled as 8 plex for two lanes on Novaseq X 10B (2×100-cycle) flow cell targeting 600M reads/samples.

The sequencing data (FASTQs) generated from the 5’ Gene Expression (GEX) libraries were processed using the 10X Genomics Cell Ranger software^48^ “cellranger multi” command with default options. Different 10X Genomics Cell Ranger versions were used to process data from different 5’ GEX library kit versions, because of differing 10X barcodes. For the single-cell 5’ v2 library kit, 10X Genomics Cell Ranger v7.2.0 was employed, while 10X Genomics Cell Ranger v8.0.1 was used for the single-cell 5’ v3 library kit. Even though this results in data from different software versions, 10X Genomics advises re-analysis with 10X Genomics CellRanger v8.0.1 is unwarranted for these library types^49^ (https://kb.10xgenomics.com/hc/en-us/articles/25011808172685-My-samples-are-analyzed-with-Cell-Ranger-v7-2-Should-I-upgrade-to-using-the-latest-Cell-Ranger-v8-0). Therefore, we used results from both software versions without any distinction. Pre-built human GRCh38 reference files, from 10X Genomics, were used for this analysis. For the GEX dataset, the sequenced reads were mapped using 10X Genomics reference version GRCh38 GEX 2020-A^50^(https://www.10xgenomics.com/support/software/cell-ranger/latest/release-notes/cr-reference-release-notes#2020-a, July 7, 2020, GENCODE v32/Ensembl 98). Metrics computed by 10X Genomics Cell Ranger were parsed and analyzed for quality control.

### Single-cell Seurat Analysis

A total of 11 single-cell samples from three individuals across three timepoints were collected for the Ax-2 space mission, and 6 samples from two individuals across three timepoints were collected for the Ax-3 mission. Sample metrics are provided in Table S1A. 5’ GEX data were analyzed using the Seurat v5 package (version 5.0.1)^51,52^. Cells identified as outliers or containing >5% mitochondrial gene content were excluded. Additionally, low-abundance genes (expressed in fewer than three cells) and potentially low-quality cells were removed from downstream analysis. The datasets contained between 200 and 5,000 genes per cell. Principal Component Analysis (PCA) was performed, and the first 20 principal components (PC1–PC20) were used to construct nearest-neighbor graphs. Louvain clustering was applied, followed by non-linear dimensionality reduction using Uniform Manifold Approximation and Projection (UMAP) for visualization. Clusters were annotated based on the expression of cell-type-specific marker genes. Visualizations included dot plots, heatmaps, violin plots, and UMAP cluster maps. Gene expression levels were reported on a log2 scale. To integrate datasets across individuals and timepoints, the Seurat v5 integration workflow was used. This approach returns a unified dimensional reduction that captures shared sources of biological variance, enabling clustering of cells in similar states across samples.

### RNA isolation

Whole blood (2.5 mL) was collected from each subject in PAXgene blood RNA tube (Fisher, Cat#23-021-01) at six time points (L-90, L-30, L-3, R + 1, R + 6 and R + 14) from Ax-2 (n=4) and at five time points (L-30, L-2, R+1, R+21 and R+60/61) from Ax-3 (n=2). The RNA was isolated as described in the GENESTAR manual^6^.

### Total RNA-Seq Library Preparation

Whole transcriptome sequencing (Total RNA-Seq) data were generated using the Illumina TruSeq Stranded Total RNA LP Globin kit (Illumina, Cat # 20020613) as previously described^53^. To monitor sample and process consistency, 1 µl of the 1:50 diluted synthetic RNA designed by the External RNA Controls Consortium (ERCC) (4456740, Thermo Fisher) was added to 1 µg of Total RNA. Additionally, the Universal Human Reference RNA (UHR) (Cat# 750500, Agilent) was processed in parallel with the RNA samples as a process control. Libraries were quantified using a Fragment Analyzer (Agilent Technologies, Inc) electrophoresis system.

### Total RNA Sequencing and Data Analysis

Sequencing was performed on the NovaSeq 6000 instrument using the S4 reagent kit (300 cycles) to generate 2×150bp paired-end reads. To achieve a sequencing depth of 50 to 100 million read-pairs per sample, between 35 and 70 RNA-Seq libraries were pooled per lane and loaded at 280 pM onto an S4 flow cell. Each lane included a 1% PhiX spike-in as an internal control to assess run quality. The RNA-Seq analysis pipeline processes raw RNA sequencing data (FASTQs), performing thorough quality control and allowing alignment to the GRCh38 reference genomes (excluding alternate contigs). It incorporates software tools including STAR v2.7.3a^54^ sequence alignment, Picard v2.22.5 for marking duplicate reads, and Samtools v1.9 for BAM to FASTQ conversion. Additionally, RSEM 169 v1.3.3^55^ is utilized for gene expression quantification, while RNA-SeQC v1.1.9^56^, 170 Qualimap2 v2.2.1^57^, and ERCCQC v1.0 generate quality metrics. The pipeline employs feature Counts v2.0.1 to derive raw gene feature counts, facilitating subsequent analysis for differential gene and isoform expression using the DESeq2 package. PCA analysis was conducted using R (version 4.4.3) and visualized using the plotly package.

### Plasma preparation

Whole blood (3 mL) was collected from each subject in one of the K2 EDTA tubes at six time points (L-90, L-30, L-3, R + 1, R + 6 and R + 14) from Ax-2 (n=4) and at five time points (L-30, L-2, R+1, R+21 and R+60/61) from Ax-3 (n=2) for plasma preparation. Plasma was isolated, and aliquots (300µL) were prepared in four matrix tubes (ThermoFisher, Cat#3741-WP1D-BR) and stored for future use, following the procedures described in the GENESTAR manual ^6^.

### Olink Explore HT assay and Sequencing

The Olink Explore HT platform is a high-throughput, multiplex proteomics technology that utilizes proximity extension assay (PEA) technology^58^. It measures over 5,400 biomarkers in a single run from minimal sample volumes with high specificity and sensitivity. This method utilizes pairs of DNA-labeled antibodies that bind specifically to target proteins, enabling quantification through DNA hybridization while minimizing cross-reactivity. It can detect proteins at sub-picogram per milliliter concentration. Plasma samples were analyzed to assess protein stability under different storage conditions using the OLINK Explore HT protocol.

The Olink Explore HT assay (Cat # 98100) was processed using automated liquid handlers (Mosquito LV genomics, Dragonfly Discovery, and Biomek FxP). Approximately 10 µL of plasma was used for the assay, and ∼5,400 proteins were assayed across 8 blocks of antibody pools. Plasma samples were then serially diluted (1:10 to 1:100,000) and loaded into two 384-well plates. Probes with unique DNA tags are divided into eight assay blocks. Undiluted plasma samples are used for blocks 1–4, and diluted plasma samples for blocks 5–8, following the standard Olink Explore HT setup followed by overnight incubation at +4°C. Only when both the antibody probe pairs bind to the target protein, the complementary oligonucleotides in proximity hybridize and are extended using a DNA polymerase. Annealed sequences are then extended, amplified, barcoded, and sequenced on the Illumina platform using NovaSeq 6000 S4 Reagent Kit v1.5 (35 cycles) to generate 24bp reads.

### Olink Data analysis

Sequence data (BCL files) were processed using Olink’s pre-processing software, *ngs2counts* (v4.7.1), to generate count data. Subsequently, Olink’s *Explore CLI* (v.2.3.1) was used to export a Parquet file containing intensity-normalized NPX values. NPX (Normalized Protein Expression) is Olink’s proprietary unit for relative protein quantification, expressed on a log2 scale.

All calculations were performed using intensity-normalized NPX values, utilizing the *OlinkAnalyze* R package (v4.2.0)^59^ (https://github.com/Olink-Proteomics/OlinkRPackage). Refer to the *OlinkAnalyze* vignette for details^60^ (https://cran.r-project.org/web/packages/OlinkAnalyze/vignettes/Vignett.html). All internal and external Olink controls were excluded. Proteins that varied ≥ 0.5 NPX across timepoints in all subjects were selected for further analysis. Linear Mixed-effects Regression (LMER) was applied to the remaining protein assays (n=3,825 for Ax-2 samples; n=4,054 for Ax-3 samples) of different timepoints: NPX∼(1|SubjectID). P-values from the regression analysis were adjusted using the Benjamini-Hochberg method, with an adjusted p-value threshold of < 0.05 considered statistically significant. This resulted in the identification of 1,112 significantly altered assays across timepoints in Ax-2 and 559 in Ax-3. Post-hoc analysis performed on the significantly changed assays determined detailed differences between timepoints, which were then visualized with PCA and heatmap plots. The R script used in this analysis is available on GitHub^47^ (https://github.com/qiaoyanw/Olink-Explore-HT-Analysis-Methods-for-TRISH_AX2_AX3-Flight/tree/main)

### Pathway Analysis Methods

To comprehensively analyze the biological pathways and processes underlying our experimental data, the ShinyGO 0.80 web-based application ^61^ was used. Differentially regulated proteins and transcripts data obtained from proteomics (OLINK), RNA-seq (bulk), and single-cell RNA-seq were analyzed to identify significantly enriched biochemical pathways. Proteins and transcripts showing statistically significant differential expression (adjusted p-value < 0.05) were selected and mapped to corresponding gene symbols. These gene symbols were then input into ShinyGO v0.80, an online gene-set enrichment tool that integrates data from multiple pathway databases, including KEGG, Reactome, and Gene Ontology (GO).

ShinyGO was used to perform pathway enrichment analysis by comparing the input gene list against background gene sets, followed by false discovery rate (FDR) correction for multiple comparisons. Pathways with FDR-adjusted p-values < 0.05 were considered significantly enriched. The results were visualized using enrichment plots and pathway networks to reveal the biological processes, molecular functions, and cellular components most affected by the proteomic and transcriptomic changes.

## Notes

### Competing Interest Statement

The authors have declared no competing interest.

