## Supplementary Tables and Figures for "Multi-Omics Characterization of Human Molecular Responses to Spaceflight Across Two Independent Missions"

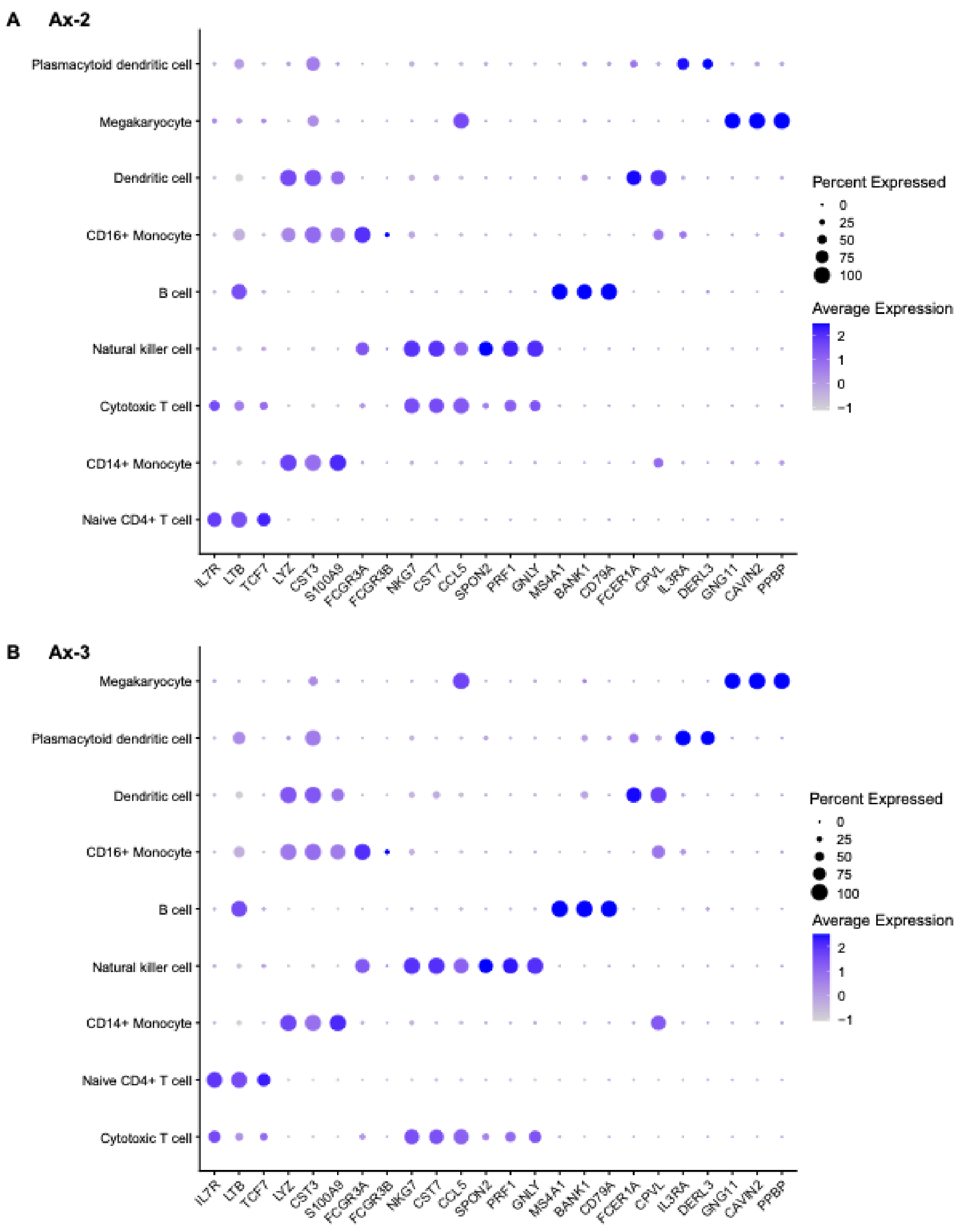


**Fig. S1: Cluster marker dot plot for Ax-2 and Ax-3 samples.** The dot plot displays the expression patterns of selected marker genes across major immune cell types for two missions: Ax-2 (top panel) and Ax-3 (bottom panel). Each row represents a cell type, and each column corresponds to a marker gene (feature). Dot size indicates the percentage of cells within a cluster expressing the gene, while color intensity reflects the average expression level (scaled). Darker shades represent higher expression, and larger dots indicate broader expression across cells. This visualization highlights distinct marker profiles that define cell identity across the two missions.

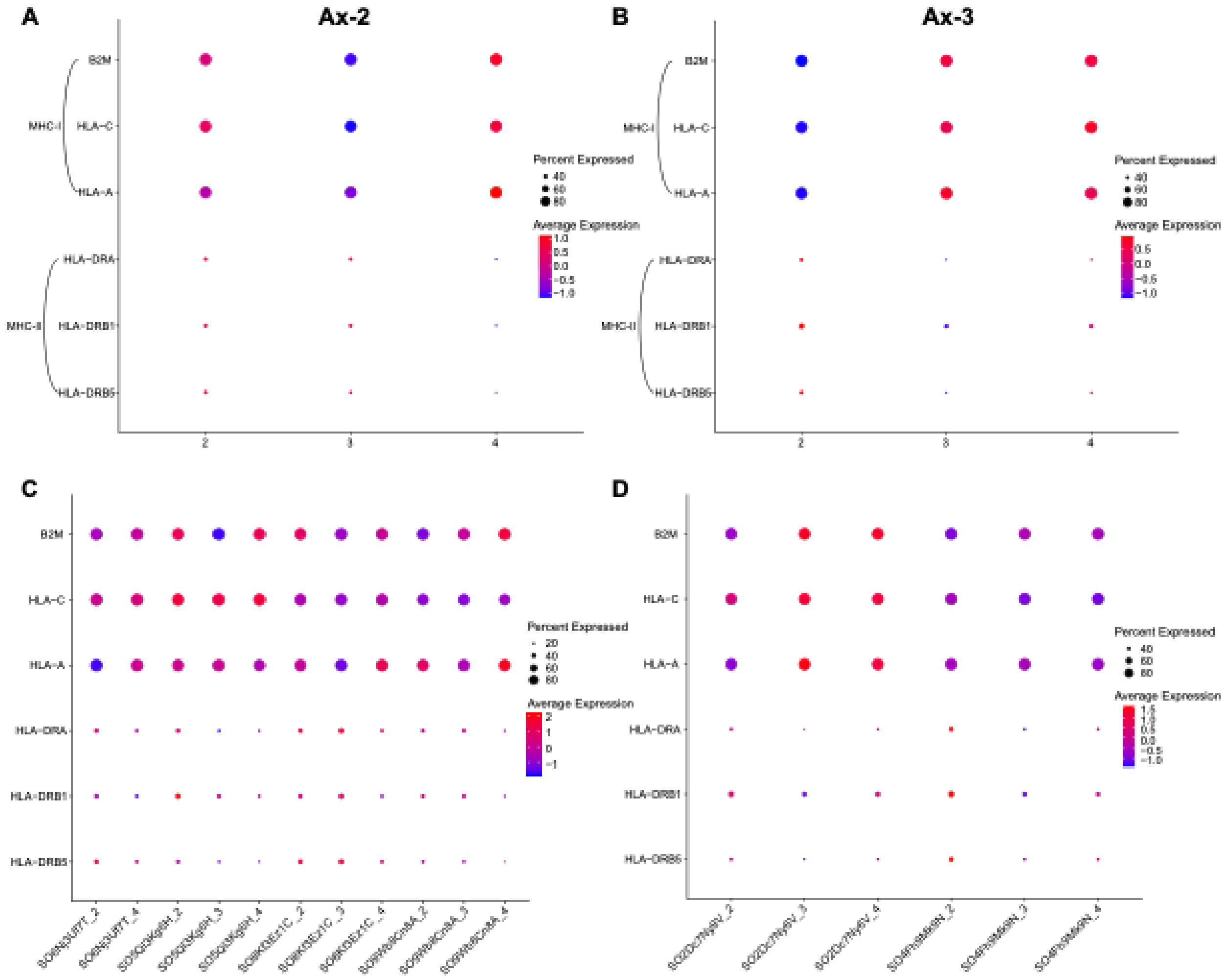


**Fig. S2: Differential expression of MHC class I and II genes.** Dot plots depict normalized expression levels of MHC-I (HLA-A, HLA-C, B2M) and MHC-II (HLA-DRA, HLA-DRB1, HLA-DRB5) genes across pre-flight and post-flight timepoints for Ax-2 and Ax-3 missions. Post-flight samples show a consistent increase in MHC-I expression and a decrease in MHC-II expression compared to pre-flight baselines.

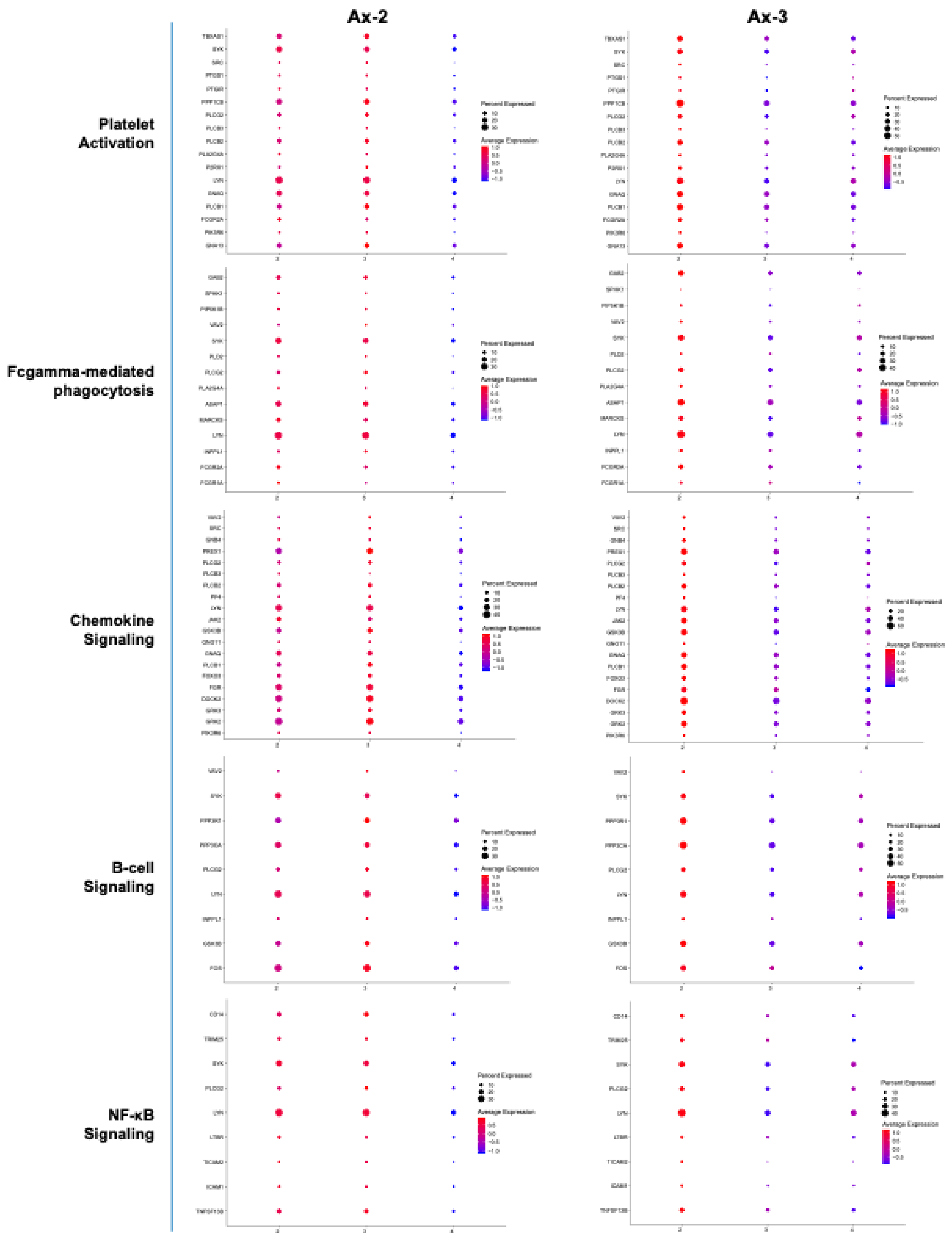


**Fig. S3A: Individual-level variation in pathway activation across Ax-2 and Ax-3 samples.** Dot plots showing changes in immune-related pathways affected in both missions, including Platelet activation, Fcγ-mediated phagocytosis, Chemokine signaling, B-cell signaling, and NF-κB signaling.

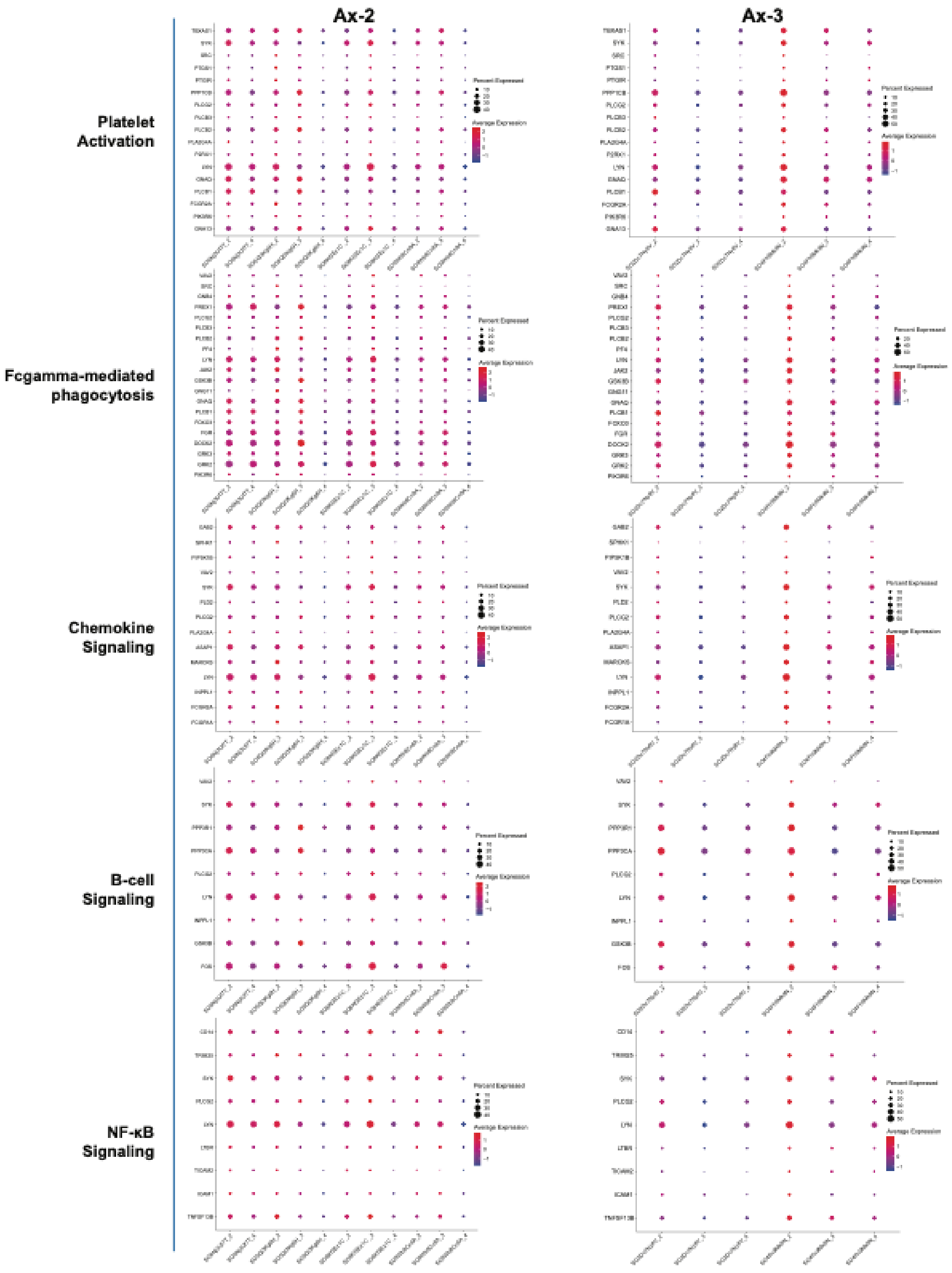


**Fig. S3B: Individual-level variation in pathway activation across Ax-2 and Ax-3 samples.** Dot plots show expression profiles of representative genes involved in Platelet activation, Chemokine signaling, Fc gamma–mediated phagocytosis, B-cell receptor and NF-kB signaling pathways across individual samples. Dot size indicates the percentage of expressing cells, and color represents the average expression level.

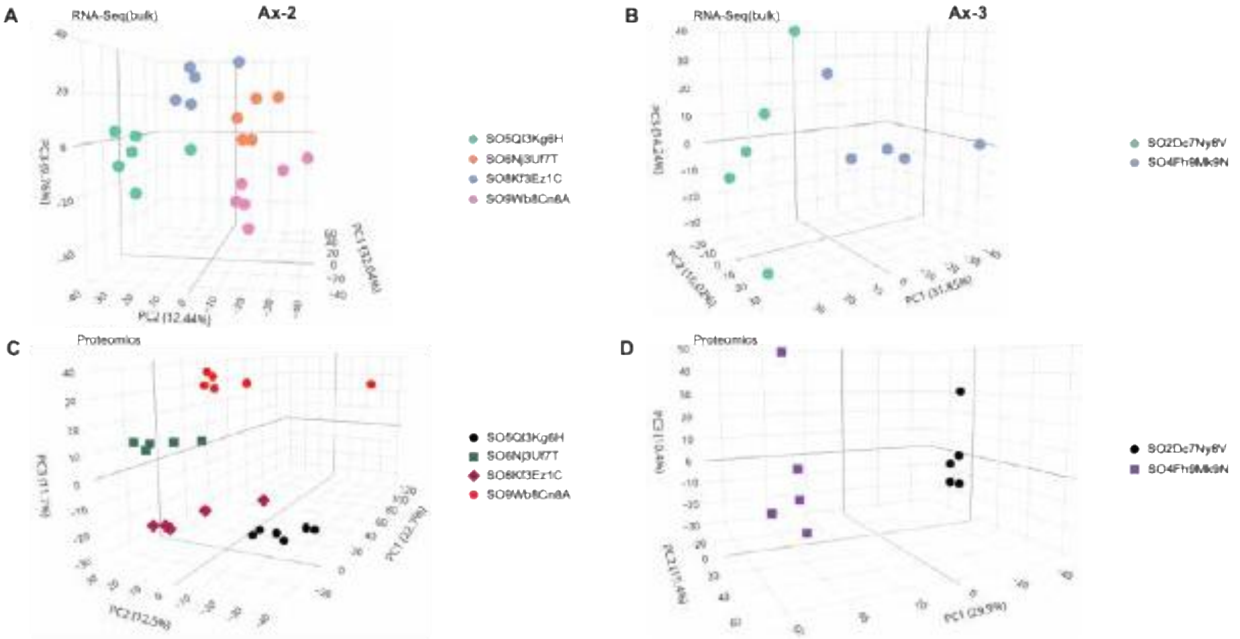


E

| Collection # | 1 | 2 | 3 | 4 | 5 | 6 | |
|---|---|---|---|---|---|---|---|
| Time of Collection | L-90 | L-30 | L-3 | R+1 | R+6 | R+14 | Total |
| Sample # | 4 | 4 | 4 | 4 | 4 | 2 | 22 |

F

| Collection # | 1 | 2 | 3 | 4 | 5 | |
|---|---|---|---|---|---|---|
| Time of Collection | L-30 | L-2 | R+1 | R+21 | R+60/61 | Total |
| Sample # | 2 | 2 | 2 | 2 | 2 | 10 |

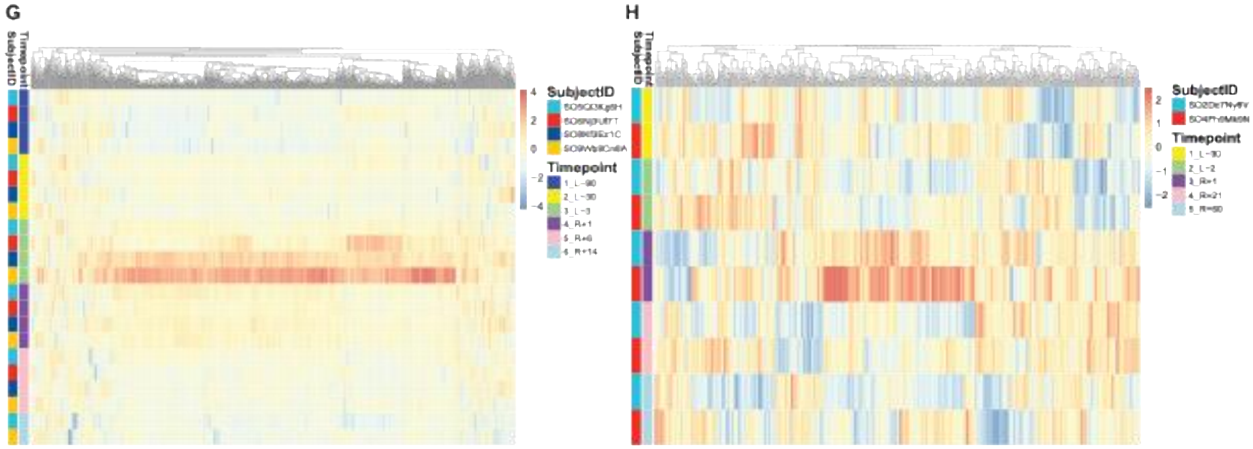


**Fig. S4**: Transcriptomic and Proteomic Profiles for Ax-2 and Ax-3 Missions.
(A, B) PCA plot of RNA-seq data showing individual-specific clustering for Ax-2 and Ax-3; (C, D) PCA plot of proteomics data showing individual-specific clustering for Ax-2 and Ax-3. Each point represents a subject, color-coded by an individual. (E, F) Sample collection overview for Ax-2 and Ax-3 missions. (G, H) Heatmaps of protein expression across timepoints for Ax-2 (L-90, L-30, L-3, R+1, R+6, R+14) and Ax-3 (L-30, L-2, R+1, R+21, R+60/61). Large vertical bars indicate timepoints; smaller bars indicate subjects. Color intensity reflects relative protein abundance.

**List of Supplementary tables**

Table S1: Single-cell sequencing matrix for Axiom-2 and Axiom-3, related to Figure 2
Table S2: List of genes commonly upregulated in Ax-2 mission L-3 vs R+1 related to Figure 3A
Table S3: List of genes commonly downregulated in Ax-2 mission L-3 vs R+1, related to Figure 3A
Table S4: List of genes commonly upregulated in Ax-3 mission L-3 vs R+1 , related to Figure 3A
Table S5: List of genes commonly downregulated in Ax-3 mission L-3 vs R+1, related to Figure 3A
Table S6: List of genes commonly downregulated in Ax-2 and AX-3 missions L-3 vs R+1, related to Figure 3A
Table S7: Sequencing Metrics: RNA-seq (bulk) libraries for 22 Ax-2 and 10 Ax-3 samples
Table S8: Molecular Responses for proteomics and transcriptomics profiles for Ax-2
Table S9: Molecular Responses for proteomics and transcriptomics profiles for Ax-3
Table S10: Analysis of non-coding RNA-Seq datasets from Ax-2
Table S11: Ax-2 Venn diagrams showing overlap among proteomics, RNA-seq (bulk), and single-cell RNA-seq
Table S12: Ax-3 Venn diagrams showing overlap among proteomics, RNA-seq (bulk), and single-cell RNA-seq
Table S13: Top 20 spaceflight responsive pathways across Ax-2 from proteomics, RNA-seq (bulk), and single-cell RNA-seq
Table S14:Top 20 spaceflight responsive pathways across Ax-3 from proteomics, RNA-seq (bulk), and single-cell RNA-seq
Table S15: Multi-Omics Analysis Data for Osteoclast Differentiation Signaling Pathway
Table S16: Multi-Omics Analysis Data for NF κB NF Signaling Pathway
Table S17: Multi-Omics Analysis Data for Coagulation, Complement, and Platelet Activation pathways
